# Diversity patterns in parasite populations capable for persistence and reinfection with a view towards the *human cytomegalovirus*

**DOI:** 10.1101/512970

**Authors:** Cornelia Pokalyuk, Irene Görzer

## Abstract

Many parasites like the *cytomegalovirus, HIV* and *Escherichia coli* are capable to persist in and reinfect its host. The evolutionary advantage (if so) of these complicated mechanisms have not been quantitatively analyzed so far. Here we take a first step by investigating a host-parasite model for which these mechanisms are driving the evolution of the parasite population. We consider two variants of the model. In one variant parasite reproduction is directed by balancing selection, in the other variant parasite reproduction is neutral. In the former scenario reinfection and persistence have been shown to sustain the maintenance of diversity in the parasite population in certain parameter regimes (Pokalyuk and Wakolbinger, 2018). Here we analyse the diversity patterns in the latter, neutral scenario. We evaluate the biological relevance of both model variants with respect to the *human cytomegalovirus* (HCMV), an ancient herpesvirus that is carried by a substantial fraction of mankind and manages to maintain a high diversity in its coding regions.

## 1 Introduction

The evolution of a host-parasite system is often interpreted in spite of an arms-race strategy, claiming that host and parasite have to adapt constantly to guarantee their own survival. However, eventually such a behaviour should lead to the ultimate extinction of the parasite or the host (while in the latter case also the parasite dies, at least if the parasite does not adapt to a new host species before). To guarantee survival the parasite rather has to evolve a strategy, which serves two conflicting forces, on the one side avoiding host impairment to prevent one’s own death and on the other side using host resources to guarantee parasite spread before the host dies (Frank, 1996). In this case an equilibrium state can be achieved at which both species coexist.

A certain amount of genetic diversity, especially in coding regions, seems to be necessary for the long-term survival of a parasite, since otherwise the host can adapt to the parasite which eventually should lead to its extinction. Random reproduction in finite populations induces a permanent decrease in diversity (Wright, 1929). Mutation is a mechanism that can restore this diversity. However, there is evidence that the bulk of the mutations happening are either neutral or deleterious for an individual (Bank et al., 2014). Hence, it should be of advantage if parasites manage to evolve mechanisms that counteract the permanent decrease in diversity in genomic regions in which diversity is beneficial.

If a parasite is capable to persist in its host, the diversity of the surrounding parasite population can be introduced in to hosts step by step by reinfection. An individual-based model for parasite evolution incorporating persistence and reinfection has been introduced in Pokalyuk and Wakolbinger, 2018. In this model the parasite population is distributed over a fixed, finite number of (infected) hosts. Two parasite types exist and each host is infected with a fixed, finite number of parasites. The composition of the parasite population may change, when parasites reproduce, when they mutate from type *A* to type *B* and vice versa, when hosts get reinfected and when an infected hosts die and is replaced by so far uninfected host, which instantly suffers from a primary infection. In the model in Pokalyuk and Wakolbinger, 2018 parasite reproduction is driven by balancing selection, i.e. there is a tendency that once a host is infected with both parasite types both parasites types coexist in this host for a long time. For this model parameter regimes have been identified, for which persistence and reinfection are effective mechanism to maintain both parasite types in the parasite population.

Here we treat the neutral counterpart, that is the model with neutral parasite reproduction. In particular, we investigate the expected diversity in the parasite population as well as the proportion of hosts expected to be infected with both parasite types.

We evaluate the biological relevance of the two scenarios at the example of the *human cytomegalovirus* (HCMV). HCMV is an old herpesvirus which is wide-spread in the human population (Cannon et al., 2010) and leads in the typical host to an asymptomatic infection (Griffiths and Baboonian, 1984; Zanghellini et al., 1999). So the HCMV population seems to have achieved a state rendering long-term coexistence with its host possible. Identifying the fundamental mechanisms allowing this long-term coexistence is also useful in the understanding the life cycle of a HCMV population in the healthy and the diseased host.

Complex mechanisms guarantee that even in the healthy host HCMV persists, reactivates and reinfects other hosts, see Collins-McMillen et al. (2018a), Collins-McMillen et al. (2018b), Goodrom (2016), and Hansen et al. (2010). With our analysis we aim to provide some basis to assess the role which persistence, reactivation and reproduction driven by balancing selection might play in ensuring long-term coexistence. For many open reading frames (ORFs) of the HCMV genome, especially coding for immune evasion strategies and cell tropism, one finds a clustering into a few genotypes. Furthermore when these ORFs have been analyzed further, one observed that hosts are often infected with several genotypes. Under neutrality - in contrast to a scenario with balancing selection - the coexistence of genotypes requires a relatively high mutation rate and a large proportion of hosts can only be infected with several genotypes, if the reinfection rate is relatively high as well. We argue that these rates might be unrealistically high for HCMV.

The manuscript is organized as follows: First we explain the model for the evolution of a parasite population with persistence and reinfection and then summarize the results from Pokalyuk and Wakolbinger, 2018, where the scenario with balancing selection has been analyzed. In the results section we analyze the neutral variant of the model and evaluate the biological relevance of the model on the basis of diversity patterns in DNA data of HCMV.

## 2 A model for parasite evolution with reinfection and persistence

We consider the following two variants of a model for the evolution of a parasite population capable for persistence and reinfection. In Variant a) parasites reproduce according to a neutral reproduction mechanism and in Variant b) parasite reproduction is affected by balancing selection. Variant b) has been analyzed in Pokalyuk and Wakolbinger, 2018.

Consider a population of parasites distributed over *M* infected hosts. Assume each host is infected with *N* parasites. We distinguish two types of parasites, type *A* and type *B*. Denote by 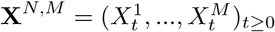 the relative frequencies of type *A* in the *M* different hosts in the course of time *t* ≥ 0.

Evolution is driven by the following four factors:

- (Parasite reproduction within hosts)
  - Variant a): Neutral parasite reproduction: The evolution of the parasite population within each host is an adaptation of the classical Moran dynamics, see Ewens, 2004, Chapter 3.4, in both variants. At reproduction a parasite splits into two and replaces with its offspring a randomly chosen other parasite. In the neutral case both types of parasites reproduce at (the same) rate *g*_*N*_. A change in frequency of type *A* only occurs if the type of the reproducing parasite differs from the type of the parasite which is replaced. Hence, if the frequency of type *A* in host *i* is *x* (for some *x* ∈ {0, 1*/N, …*, (*N* − 1)*/N*, 1}) a jump due to parasite reproduction to 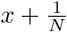 occurs at rate *g*_*N*_ *Nx*(1 *- x*) and to 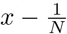 at rate *g*_*N*_ *Nx*(1 *- x*).
  - Variant b): Parasite reproduction with balancing selection: We assume there exists some equilibrium frequency *η* (with *η* ∈ (0, 1)), at which type *A* and *B* are optimally balanced in a host. If the frequency of type *A* in host *i* is *x* each parasite of type *A* reproduces in this host at rate *g*_*N*_ (1 + *s*_*N*_ (*η − x*)) and parasites of type *B* at rate *g*_*N*_ (1 + *s*_*N*_ (1 − *η* − (1 − *x*))) = *g*_*N*_ (1 − *s*_*N*_ (*η* − *x*)), where *s*_*N*_ > 0 denotes the selection strength. That is the rate of reproduction of *A*-type parasites is larger than that of *B*-type parasites, if the frequency of type *A* is below the equilibrium frequency *η*, at which type *A* and type *B* are optimally balanced, and vice versa if the frequency is above the optimal frequency. If a parasite reproduces it doubles and replaces with its progeny a randomly chosen parasite in this host. A change in frequency only occurs if the type of the reproducing parasite differs from the type of the parasite which is replaced. Hence, if the frequency of type *A* in host *i* is *x* (for some *x* ∈ {0, 1*/N, …*, (*N* − 1)*/N*, 1}) jumps to 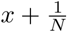 occur due to parasite reproduction at rate *g*_*N*_ (1 + *s*_*N*_ (1 + *s*_*N*_ (*η − x*)))*Nx*(1 *- x*) and to 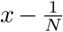 at rate *g*_*N*_ (1 + *s*_*N*_ (*x − η*))*Nx*(1 *- x*).
- (Host replacement) Hosts die at rate 1. Whenever a host dies it is replaced by a so far uninfected host, which instantly suffers from primary infection. At primary infection the host is infected with a single type chosen randomly according to the type frequencies in the transmitting host. This leads to a jump to frequency 0 or 1 in the primary infected host. Hence, if the frequency of type *A* in hosts 1, *…, M* is *x*_1_, *…, x*_*M*_ in host *i* jumps to state 1 occur at rate 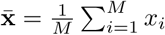 and to state 0 at rate 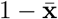.
- (Reinfection) In addition to parasite reproduction, hosts get reinfected at rate *r*_*N*_. At reinfection a single parasite, which is chosen randomly in the reinfected host, is replaced by a randomly chosen parasite of the reinfecting host. Consequently, changes in frequencies are only observed, if the replaced parasite has another type as the transmitted parasite. Hence, if the frequency of type *A* in hosts 1, *…, M* is *x*_1_, *…, x*_*M*_ in host *i* jumps due to reinfection to 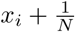 occur at rate 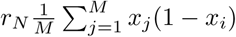 and at rate 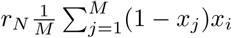 to 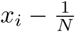.
- (Mutation) Each parasite mutates at rate *μ*_*N*_ to type *A* and at the same rate to type *B*.

To summarize jumps from state

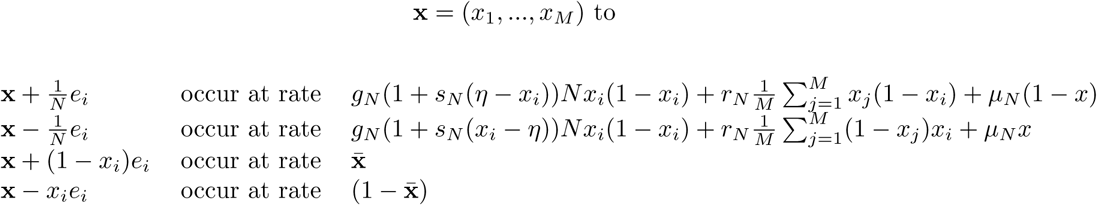

for *i* = 1, *…, M* with *x*_i_ ∊ {0,1/*N*, … 1}, 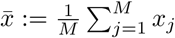 and *e*_*i*_ = (0, *…*, 1, *…*, 0) the *i*-th unit vector of length *M*.

### 2.1 Parasite reproduction with balancing selection

Variant b) of the model has been analyzed in detail in Pokalyuk and Wakolbinger, 2018. We give here the upshots of the results obtained therein concerning the diversity patterns and maintenance of diversity.

Assume the host population size *M* as well as the parasite population size *N* are large and the following conditions are fulfilled:

Assumptions:

1. (moderate selection)

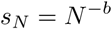

for some 0 < *b* < 1.
2. (frequent reinfection)

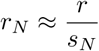

for some *r* > 0.
3. (fast parasite reproduction)

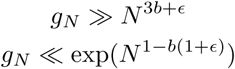
4. (rare mutation)

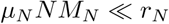

Then for large *N* host type frequencies are concentrated around 0, *η* and 1. Denote by *v*^*l*^ the proportion of hosts which frequencies are concentrated around *l* for *l* = 0, *η*, 1. Then for large *N* the evolution of the frequencies (*v*^0^, *v*^*η*^, *v*^1^) is driven by the dynamical system

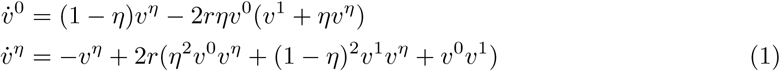

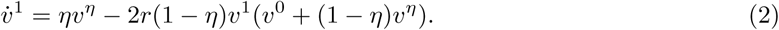

For

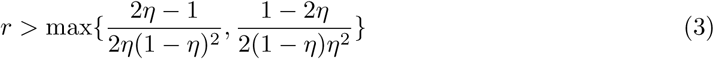

the dynamical system **v** = (*v*^0^, *v*^*η*^, *v*^1^) has a unique stable equilibrium in the point **u** = (*u*^0^, *u*^*η*^, *u*^1^) with

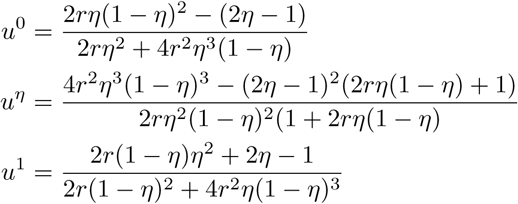

(and saddle points in (1, 0, 0) and (0, 0, 1)).

The stability of the dynamical system is a key ingredient to show, that under Assumptions 1-4 and 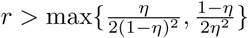 (which is a slightly stronger assumption on *r* than (3)) most of the time both types are present in the parasite population at non-trivial frequencies. Indeed in Theorem 3 in Pokalyuk and Wakolbinger, 2018 it is shown that under these assumptions the time until an initially monomorphic parasite population (that is a parasite population for which all parasites are either of type *A* or all parasites are of type *B*) turns into a parasite population with host type frequency proportions close to the equilibrium state **u** is of the order 1/(*μ*_*N*_ *MNs*_*N*_) + *M*^*γ*^ for any *γ* > 0 and large *N* and *M* and vice-versa the time until a parasite population with host type frequency proportions close to the equilibrium state **u** is turned into a monomorphic population is exponentially large in *M*, more precisely this time is bounded from below by exp(*M* ^1*-γ*^) for any *γ* > 0. Hence, if 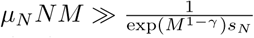 (which is a very weak lower bound on the mutation rate) most of the time both parasite types coexist in the parasite population.

In Figure 2 a simulation of the model is depicted. Except for the host population size, which has been chosen for feasibility of the simulations, the parameter values could be realistic for HCMV.

**Figure 1:**
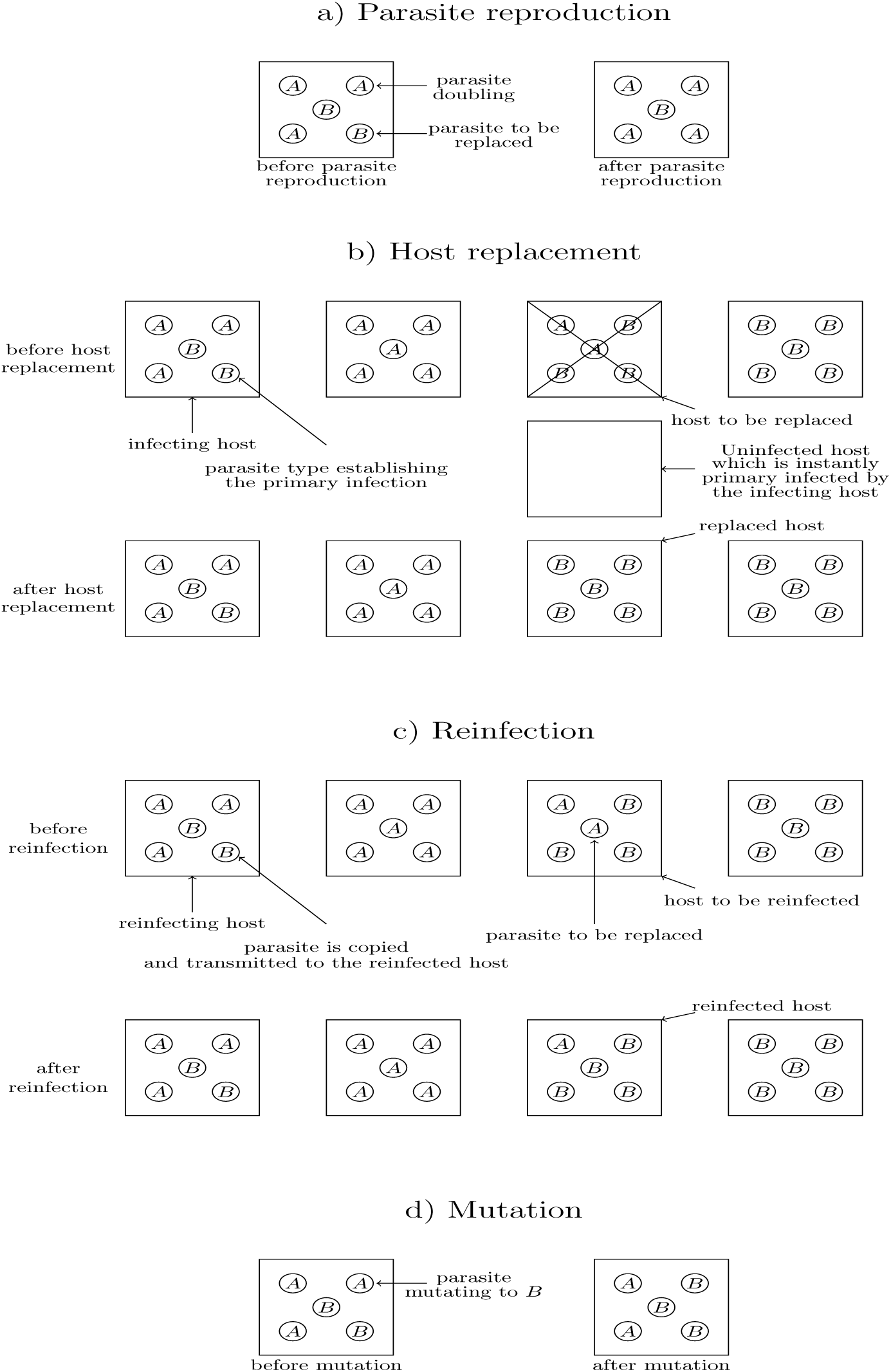
The parasite population evolves according to the four mechanisms a) parasite reproduction, b) host replacement, c) reinfection and d) mutation, see also the Definition in Section 2: a) Parasites reproduce within hosts by cloning a parasite and simultaneously substituting a parasite. The rates at which parasites reproduce depend on the type of the parasite and the frequency of the parasite type in the host. b) Hosts die at a constant rate. When a host dies it is instantly replaced by a so far uninfected host, which is directly infected by a randomly chosen host. At primary infection only a single parasite type is transmitted. c) Hosts reinfect other (randomly chosen) hosts at a constant rate. At reinfection a single parasite is cloned and transmitted to the reinfected host. The transmitted parasite replaces a randomly chosen parasite in the reinfected host. d) Each parasite mutates at a (small) constant rate to type *A* and type *B*.

**Figure 2:**
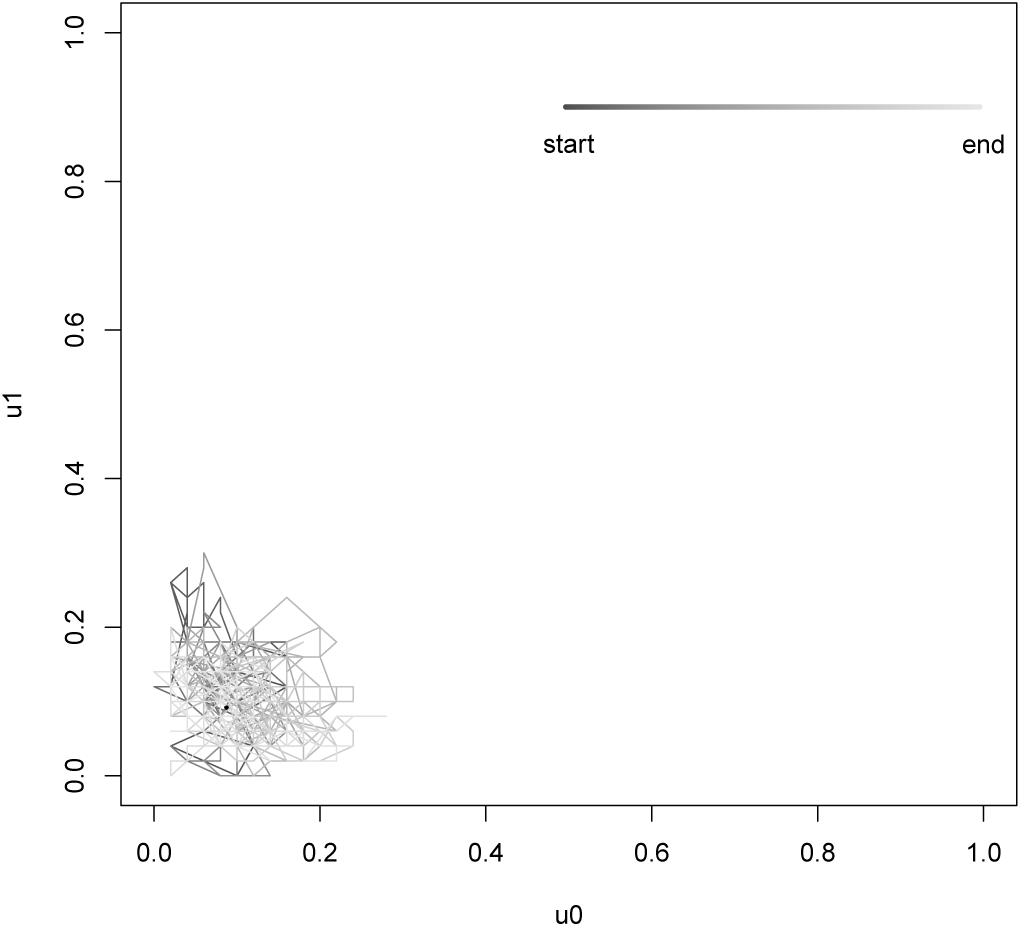
A simulation of the model with balancing selection for *η* = 0.5, *r* = 9, *g*_*N*_ = 1500, *M*_*N*_ = 50, *N* = 1000, *b* = 0.25 for 10^9^ events, that is roughly 13000 host time units. Every 10^3^ events the proportion of hosts with a frequency of type *A* below *s*_*N*_ is plotted against the proportion of hosts with a frequency of type *A* above *s*_*N*_. The thick black dot indicates the location of the equilibrium point **u**.

## 3 Results

### 3.1 Diversity patterns under neutrality

In this section we analyze the neutral setting. We consider two cases. In the first case the two parasite types are constantly generated by mutation, in the second case the two types are assumed to be present in the population due to standing genetic variation, i.e. due to an equilibrium of genetic drift and mutation. We will evaluate the role of the neutral scenario in the case of HCMV by assessing genotyping data of HCMV in Section 3.2.3 and Section 3.2.4. We make the following assumptions:

- In both cases we assume as before that the parasite populations within hosts as well as the host population is large, that is we are interested in the case when *N* and *M* is large. In precise mathematical terms we will consider the limiting case *N* → ∞ and *M* = *M*_*N*_ → ∞ for *N* → ∞.
- As before parasite reproduction is frequent. More precisely, we assume

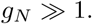
- We consider a regime in which the reinfection rate is at most of the same order as the reproduction rate, i.e. either

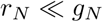

or

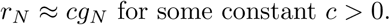

The biological motivation behind this assumption is that prior to the reinfection of a host in the transmitting host the (non-persistent) parasite population has to grow to some certain (large) size. This presumably requires also (many) reproduction events within the persistent parasite population. In particular, this assumption should apply, if the virus has a low virulence, for further discussion see Section 3.2.

#### 3.1.1 Diversity due to mutation

Consider the model from Section 2 with neutral parasite reproduction and assume that the just stated assumptions are fulfilled.

We show next, see also Figure 3, that the length of the total genealogy of a sample of size *n* drawn randomly from the parasite population is at most of order log(*n*) min {*M, NM/r*_*N*_} with high probability. That is if *N* is large the length of the total genealogy can be estimated from above by *c*_1_*log*(*n*) min {*M, NM/r*_*N*_}, for some appropriate constant *c*_1_ > 0, with a probability that is approximately 1.

**Figure 3:**
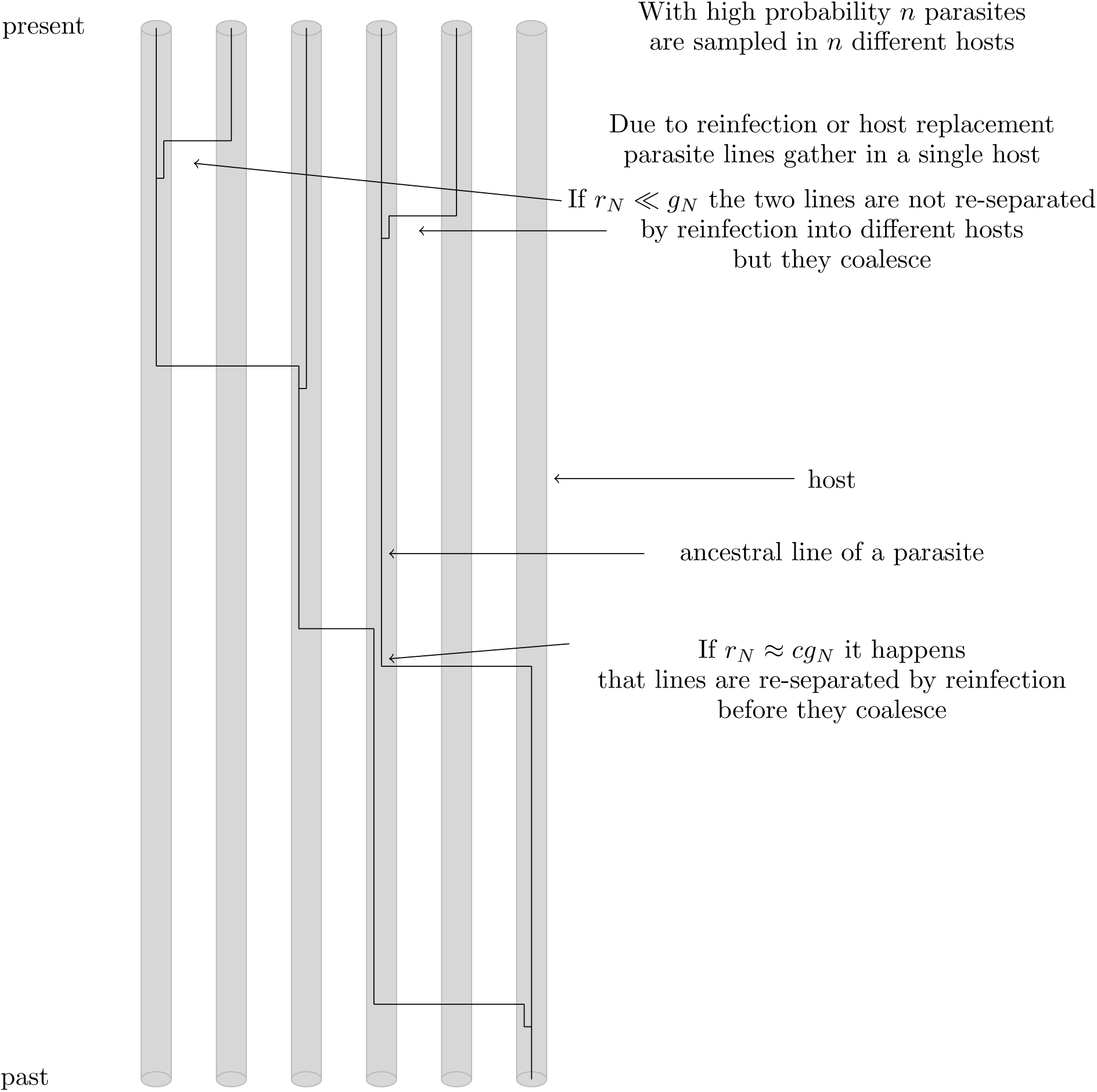
Sketch of the arguments leading to the estimate that the total length of the genealogy is of order *O*(log(*n*) min {*M, NM/r*_*N*_}). In the figure *n* = 5 parasites are sampled randomly from the total parasite population in the present and the ancestral lines of the parasite are followed backwards until all lines have coalesced.

Draw a random sample of size *n* from the parasite population and follow back its ancestry. As the parasites are sampled randomly and the host population is large the parasites are drawn with high probability from *n* different hosts.

Since parasite reproduction is restricted to hosts, to join ancestry parasites have to infect a common host first. This may happen via reinfection or host replacement (i.e. primary infection).

Each pair of parasite lines is gathered in a single host due to reinfection at rate 2*r*_*N*_ /(*NM*), because each of the two lines reinfects other hosts at rate *r*_*N*_ */N* and the probability that the host infected by the other parasite line is involved in the reinfection event is 1*/M*. Similarly each pair of lines is gathered in a single host due to host replacement at rate 2*/M*, because each host is replaced at rate 1 and so each parasite line is transfered backwards in time to is hit by a host replacement event at rate 1 and the probability that the host infecting is also the one infected by the other parasite line is 1*/M*.

Consequently the rate at which two out of *n* lines gather in a single host due to reinfection or host replacement is 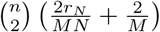.

Once two lines infect the same host, they coalesce with some positive probability (due to parasite reproduction or host replacement) before a reinfection event can separate lines again into different hosts. Indeed, each host is hit by an host replacement event at rate 1 and each pair of lines coalesces at a parasite reproduction event at rate 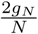. Since by assumption *r*_*N*_ ≪ *g*_*N*_ or *r*_*N*_ ≈ *cg*_*N*_ for some *c* > 0 and each line is hit by a reinfection event at rate *r*_*N*_ */N*, the probability that the two lines coalesce before they are hit by reinfection event is strictly larger than 0.

If two lines coalesce, the time till coalescence is exponentially distributed with rate 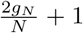. Since 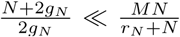 this time is negligible in comparison to the time needed for two lines to gather in a host due to reinfection or host replacement.

Consequently, the time scale of the genealogy is min{*M, MN/r*_*N*_ }. If time is accelerated by this factor, the genealogy is asymptotically given by a Kingman coalescent. If *r*_*N*_ ≪ *g*_*N*_ this is immediate, because on the appropriate time scale the time until two lines located in different hosts gather in a single host is asymptotically exponentially distributed and the time until two lines located in the same host coalescence is negligible for large *N*. If *r*_*N*_ ≈ *cg*_*N*_, for some *c* > 0, an asymptotically geometric number of trials is necessary until two lines after gathering in a single host also will coalesce in a host. The waiting time for two lines to gather in a single host is asymptotically exponentially distributed (on the Kingman time scale). As a geometric number of exponentially distributed waiting times is also exponentially distributed ^1^ the genealogy converges in the case *r*_*N*_ ∼ *cg*_*N*_ for some *c* > 0 to a Kingman coalescent as well. For a Kingman coalescent of size *n* it is well known, that the total tree length has expectation ∼ 2 log(*n*) and a finite variance, see Wakeley, 2009, Chapter 3.

Hence, by Chebychev’s inequality, the length of the total genealogy is of order

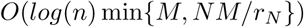

with high probability.

### Remark 1

(Transmission of multiple types at primary infection). *In both variants of our model we assume that at primary infection only a single type is transmitted. If the primary transmission process lasts for a long time, like in the case of HCMV for which many hosts are primary infected by their mothers via breast feeding, multiple types could be transmitted, see also the point “primary infection with a single type” in Section 3.2.1.*

*In a more general model of primary infection host replacement can just have a weaker effect and therefore in the general case the length of the total genealogy is of order O*(*log*(*n*)*NM/r*_*N*_) *and not of order O*(log(*n*) min{*M, NM/r*_*N*_}).

#### 3.1.2 Diversity due to standing genetic variation

A certain locus might also be diverse due to standing genetic variation. Assume the frequency of type *A* in the parasite population is 0 < *x* < 1 and *μ*_*N*_ ≪ 1. We are interested in the distribution of the frequency *W*^*N*^ of type *A* in a randomly chosen host. We distinguish the following cases:

a. Assume *r*_*N*_ ≪ *g*_*N*_. Then *W*^*N*^ converges (in distribution) for *N* → ∞ to a Bernoulli-distributed random variable with parameter *x, Ber*(*x*) for short. That is for large *N* the proportion of hosts, for which the frequency of type *A* is close to 0, is almost *x* and the proportion of hosts, for which the frequency of type *A* is close to 1, is almost 1 *- x*.
b. Assume *r*_*N*_ ≈ *cg*_*N*_ for some *c* > 0 and assume 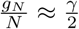 and 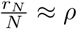 for some 0 ≤ *γ, ρ* ≤ ∞.
  i. If *ρ* = 0, then *W*^*N*^ converges (in distribution) to a *Ber*(*x*)-distributed random variable.
  ii. If *ρ* = ∞, then *W*^*N*^ converges (in distribution) to a *Beta*(*xc*, (1 − *x*)*c*) distribution. That is for large *N* the proportion of hosts for which the frequency of type *A* lies between 0 < *q*_1_ < *q*_2_ < 1 is 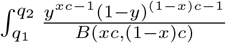 with 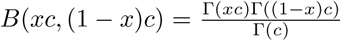 und Γ denotes the Gamma function.
  iii. If 0 < *ρ, γ* < ∞, *W*^*N*^ converges (in distribution) to a mixture of Beta distributions. This mixture has more mass close to the boundaries 0 and 1 than the corresponding Beta distribution in case (ii) (with the same parameter *c*).

To see the above statements consider a random sample of parasites of size *n* taken from a randomly chosen host and follow back its ancestry, see also Figure 4. To arrive at the frequency of type *A* in the chosen host, we need 1 ≪ *n*, but we are free to choose *n* ≪ *N*.

**Figure 4:**
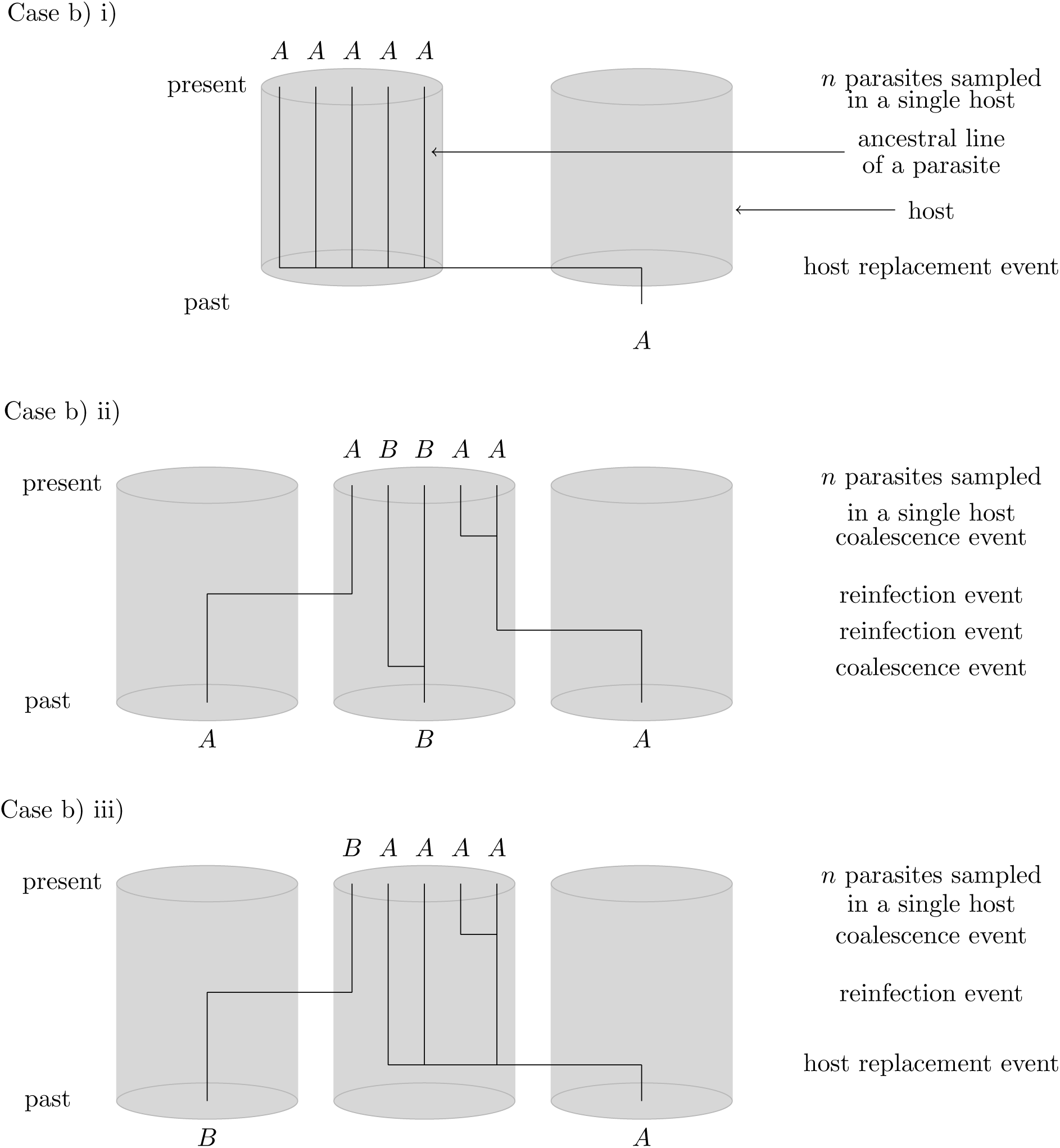
Example for genealogies in the cases b) i) - iii). *n* = 5 parasites are sampled in a single host and the genealogy of the sample is followed backwards until each host is infected at most by a single parasite line of the sample. Then the types of these lines are specified by randomly sampling both types according to their frequencies at that time point in the past. Thereafter types are propagated further along the genealogy till the present when parasites are sampled.

By the same arguments as in Section 3.1.1 the *n* lines coalescence (due to parasite reproduction) at rate *n*(*n* - 1)*g*_*N*_ */N* and the *n* lines are hit by reinfection events at rate *nr*_*N*_ */N.* Finally hosts are replaced at rate 1, in this case all lines are involved simultaneously. Consequently the relationship of coalescence: reinfection: host replacement is

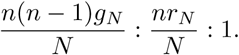

Case a): In a standard Kingman coalescent with a pair coalescence rate of 1 the time until *n* lines coalesce has expectation 2(1 − 1*/n*) and variance 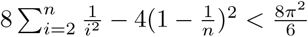, see Wakeley, 2009. In our setting pairs of parasites coalesce at rate 2*g*_*N*_ */N* due to parasite reproduction and all lines coalesce due to host replacement at rate 1. Hence, in expectation all lines coalesce till time ≈ 4*N/g*_*N*_, if *g*_*N*_ ≫ *N*, and accordingly the variance is at most proportional to (*N/g*_*N*_)^2^, and otherwise all lines coalesce within a time frame of order 1. Hence, the total length of the genealogy is of order *n* and so with high probability no mutations fall on the total genealogy, as we assumed that *μ* ≪ 1.

Every parasite is hit by a reinfection event at rate *r*_*N*_ */N*. By assumption *r*_*N*_ ≪ *g*_*N*_ hence, by Chebychev’s inequality with high probability all lines coalesce (due to parasite reproduction or host replacement) before a reinfection kicks in.

The type of the MRCA (most recent common ancestor) is a random sample from the parasite population. Host replacement occurs at rate 1 per host and the time to the MRCA of the total host population is of order *M*_*N*_. Hence, the time to the MRCA within a host is small in comparison to the MRCA of the total parasite population and therefore the frequency of type *A* in the total parasite population is still approximately *x* when the MRCA of the sample is reached. Hence the probability that the MRCA of the sample is of type *A* is approximately *x*. As no mutations fall on the genealogy with high probability the type of the MRCA is propagated further to all parasites in the sample at present.

Case b): Parasite reproduction and reinfection act here on the same time scale. The diversity pattern depends on the relationship of the rate of parasite reproduction/reinfection and the host replacement rate. The total length of the genealogy is in all three cases of order *n* (or less). Hence as in Case a) mutation can be ignored.

In Case b)i) *ρ* = 0, i.e. *r*_*N*_ ≪ *N* as well as *g*_*N*_ ≪ *N*. Consequently, a host replacement event hits the ancestral lines of the *n* parasites much earlier than host replacement and reinfection. At a host replacement event all lines coalesce into a single line. Consequently, as in Case a) with probability *x* the sample consists only type *A* parasites, with probability 1 − *x* only of type *B* parasites.

In Case b)ii), host replacement is negligible for large *N*. The relationship of coalescence: reinfection reads

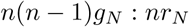

which simplifies to

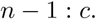

Typically in this scenario some lines are hit by reinfection events before the MRCA of the *n* lines is reached. If an ancestral lineage is hit by a reinfection event the line is followed further in a randomly chosen host. If *k* (for some *k* ∈ {1, *…, n*}) lines are hit by a reinfection event, all these lines are followed up backwards in time in *k* different hosts with high probability as *M* is by assumption large. Furthermore, since the MRCA of the remaining lines is found within a time frame of order 1, no recognizable change of the frequency of type *A* occurs in the limit *N* → ∞. Hence, at the time point in the past when each host is infected at most with a single ancestral line of the sample, each of these lines is, independently of the others, with probability *x* of type *A* and with probability 1 − *x* of type *B*. As the total length of the genealogy is of order *n* This scenario can also be interpreted as the genealogy of a Wright-Fisher diffusion with parent independent mutation to type *A* at rate *xc* and to type *B* at rate (1 − *x*)*c*. In this case the distribution of the frequency of type *A* is given by a Beta-distribution with parameters *cx* and (1 − *c*)*x*, see Ewens, 2004, Chapter 5.6, or Chapter 1.3.4 in Birkner, 2005. Indeed, forward in time the genealogy of the sample of parasites can be generated via a Yule process with immigration, see Chapter 1.3.4 in Birkner, 2005 or for similar results Donnelly and Kurtz, 1996. Yule-trees of type *A* (of type *B*, respectively) immigrate at rate *xc* (at rate (1 *- x*)*c*) and each branch splits at rate 1. Let 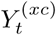 and 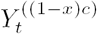 be the number of branches at time *t* in the two Yule processes with immigration started at time 0. 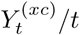 converges to a Gamma-distributed random variable with parameter *cx* and 1, see the proof of Theorem 4 in Birkner, 2005. Due to the properties of sums and quotients of independent Gamma-distributed random variables, for *n* → ∞ the proportion of type *A* branches 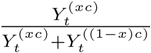 is Beta(*xc*, (1 *- x*)*c*) distributed.

In Case b)iii) all three forces host replacement, reinfection and parasite reproduction are acting on the same time scale. The relationship of parasite reproduction: reinfection: host replacement reads

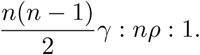

Coalescence and reinfection alone create the Beta distributions as in case b) ii), but host replacement ends up the genealogy earlier giving reinfection less chance to introduce diversity.

Consequently, more lines are in general of the same type and the distributions of the frequencies have a heavier mass close to 0 and 1.

Indeed, we can construct the genealogy of a sample of size *n* as follows:

Consider first a coalescent started with *n* lines which lines are killed at rate *ρ* and with pairs of lines coalescing at rate *γ*. Let *K*^*n*^ be the (random) number of lines left in such a coalescent when a clock with an exponentially distributed waiting time with parameter 1 rings. These *K*^*n*^ lines are left, when the host replacement event occurs. Hence, these lines must have all the same type. With probability *x* they are of type *A* and with probability 1 − *x* they are of type *B*. Furthermore, since at each event (coalescence or reinfection) the number of lines followed up further decreases by 1, the probability that the *i*-th event is a coalescence and reinfection event, resp., is independent of the event *K*^*n*^ = *k*, for 1 ≤ *i* ≤ *k* and *k* ≤ *n*. Hence, to arrive at the type distribution of a sample of size *n* one can also use a backward construction as follows: Start with *K*^*n*^ with *k* lines according to the backwards construction and type all lines with probability *x* with type *A* and otherwise with type *B*. Now add lines until *n* lines are in the sample as follows: Lines branch at rate 1 and lines of type *A* immigrate at rate *cx* and lines of type *B* immigrate at rate *c*(1 − *x*).

Let *K*∞ be the (random) number of lines left, if the coalescent above is started with ∞-ly many lines and construct in the same manner as in the paragraph before. Then as in case b)ii) the limiting distribution of the proportion of type *A* branches is with probability *x Beta*(*K*∞ + *cx, c*(1 − *x*)) and with probability 1 − *x Beta*(*cx, K*∞ + *c*(1 − *x*)). This can be seen as follows: Let *Y*_*t*_ be the number of branches in a Yule process, for which every branch splits at rate 1, at time *t*. Then *Y*_*t*_*/e*^*t*^ converges for *t* → ∞ to an exponentially distributed random variable with parameter 1, see Chapter 1.3.4, Lemma 2 in Birkner, 2005. Furthermore, let 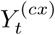 the number of branches in a Yule process with immigration, for which every branch splits at rate 1 and branches immigrate at rate *cx*, at time *t*. Then 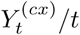 converges to a Gamma-distributed random variable with parameter *cx* and 1 as in Case b)ii).

If *K*∞ = *K* and initially all lines are of type *A* the proportion of lines of type *A* is

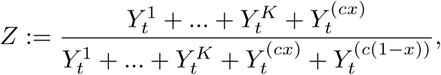

since lines of type *A* immigrate at rate *cx* and lines of type *B* immigrate at rate *c*(1 − *x*). *Z* converges for *t* → ∞ to Beta(*K* + *cx*, (1 − *x*)*c*), due to the properties of sums and quotients of independent Gamma-distributed random variables. In the same manner the proportion of lines of type *B* is obtained.

### Remark 2

(Transmission of multiple types at primary infection). *As in Section 3.1.1 we also want to discuss more general models of primary infection. In Section 3.2.2 we will apply the results in the scenario b) ii), because under this scenario the highest frequency of hosts infected with both genotypes is expected.*

The asymptotics in this case are not changed, if at primary infection also more than one type can be transmitted. Indeed, in this case host replacement events (at which primary infection occurs) are not relevant, since the ancestry is (asymptotically) already determined before a host replacement occurs.

In case a) host replacement may occur before the MRCA of the sample is found. However if multiple virions may be found the persistent population, from a backwards view this just means that these lines have to be followed further in another, but identical host. Since in case a) reinfection is negligible all lines will coalesce before reinfection can separate the lines into different hosts. Consequently, in case a) host type frequencies are still Bernoulli-distributed if at primary infection multiple types can be transmitted.

In case b) i) lines merge predominantely due to host replacement, if at primary infection only a single genotype is transmitted. If at host replacement only a small number of virions found the persistent population after a few host replacement events all lines merge into one, so that reinfection cannot separate lines into different hosts and we end up at a Bernoulli-distribution again. If the effective number of founding virions at primary infection is larger, reinfection might shift the distribution of host type frequencies towards a distribution with some mass at intermediate host type frequencies, because backwards in time reinfection could separate whole families of virions which coalesced already at host replacement events closer to the present time.

In case b) iii) the effect of host replacement is a shift of host type frequencies towards the boundaries 0 and 1. In a general model of primary infection this effect is weakened.

### 3.2 Biological relevance

The model is mainly motivated by observations made for the *human cytomegalovirus*. However, the model described above and the parameter regimes we consider in this study might be also relevant for other parasites. In the following subsection we gather some information about HCMV from the literature, substantiating the biological relevance of the model. In particular, we discuss which parameter regimes could be relevant for HCMV. These parameter estimates will then be used in Subsections 3.2.3 and 3.2.4 to assess the relevance of the neutral scenario for HCMV on the basis of genotyping data.

#### 3.2.1 Applicability to HCMV

HCMV is a herpesvirus that persists in its hosts after primary infection as well as after reinfection, that is once infected a host carries a HCMV population established at possibly several infection events till the end of its life. During persistence the viral population switches between phases of reactivation and latency. In this manner the virus manages to hide from the host immune system in phases of latency, but can spread at phases of reactivation. Complex mechanisms control the switches, see Dupont and Reeves, 2016.

In DNA regions encoding proteins important for cell-to-cell virus transmission and host cell entry as well as encoding strategies of immune evasion and immune modulation ^2^ and regions encoding proteins permitting the HCMV cell tropism ^3^ one finds a clustering into a few phylogenetically distinct genotypes, see Puchhamer-Stöckl and Görzer, 2011 for a review. These genotypes have been shown to evoke different immune responses of the host (for UL75 (encoding glycoprotein gH), see Urban et al., 1992 and for UL73 (encoding glycoprotein gN), see Burkhardt et al., 2009; Park, 2021 and a modified protein function (for UL55 (encoding for glycoprotein gB), see Stangherlin et al., 2017). Considering single open reading frames (ORFs) there are non-trivial proportions of hosts infected only with a single genotype and non-trivial proportions of hosts infected with multiple types, see e.g. Paradowska et al., 2014 for the ORF UL75, Novak et al., 2008; Rycel et al., 2015; Bale et al., 1996 for UL55, Murthy et al., 2011 for UL55, UL144, UL146 and UL09. When an infected host is reinfected with a virus carrying a genotype that is absent in this host, this genotype may evade preexisting immune response to establish persistence and enhance reactivation of the virus due to a missing immune response on the host’s part Burkhardt et al., 2009. Furthermore, a certain diversity present in an infected host may act as a permanent stimulator for reactivation, as a diverse viral population might be more difficult to be controlled than a monotone one. With our model and its analysis we aim to give some insights how these infection patterns with hosts infected with a single genotype and hosts infected with several genotypes may arise in a parasite population.

Our model is designed to be in line with the observed evolution of HCMV. Next we discuss the model design with respect to the HCMV. In the literature most studies investigated HCMV infections in transplantation patients and congenitally infected children in whom viral reproduction is often more enhanced than in the healthy immunocompetent host. On an evolutionary time scale these patients are probably of low relevance, therefore here we primarily refer to the literature analyzing the infection dynamics in healthy hosts.

- (Carrying capacity of within host parasite populations) We assume in our model that there exists some carrying capacity that the viral population maintains in the host also at phases of latency. For simplicity we assume, that this carrying capacity does not change over time and does not differ between hosts. It is currently unknown, if these model assumptions apply for HCMV. The persistent population may be increasing as long as the immune system cannot control HCMV sufficiently (the controllability of HCMV depends on the age of the host: in young, infected children (with and without symptoms) high HCMV DNA loads can be detected for more than a year, Adler, 1991a, in adults the virus sporadically reactivates, see Boven et al., 2017; Huang et al., 2018). It is possible that the viral population size is decreasing over time when the virus does not manage to reactivate within a host or reinfect a host sufficiently often, since cells in which the virus persists are replaced. The host immune response on HCMV certainly differs between hosts. This could also effect the carrying capacities. However, at least the number of latently infected mononuclear cells ^4^ did not vary a lot in eight different, presumably immunocompetent hosts, see Slobedman and Mocarski, 1999. Albeit the number of examined patients is rather small, this finding can be taken as an indication for rather uniform carrying capacities among hosts. The parameter regimes which could be relevant for the size of within host parasite populations are discussed separately under the point “parameter regimes”.
- (well-mixed host population and well-mixed parasite populations within hosts) To the best of our knowledge, so far neither estimates of the host population size, nor an analysis of the host population structure is available. Only a few studies analyze the geographic structure of the HCMV population. In Sijmons et al., 2015 132 full genome sequences have been analyzed with 130 sequences from Europe, one from the USA and one from South Korea. The estimated phylogeny exhibits a star-like form, the two sequences outside of Europe did not cluster separately. Pignatelli et al., 2003 analyzed the worldwide distribution for UL 73 (for samples from Europe, China, Australia and Northern America) and Bradley et al., 2008 for UL146 und UL 139 (for samples form Africa, Asia, Australia and Europe). No significant differences were detected in the genotype distributions. Therefore the simplistic assumption of a panmictic, i.e. well-mixed, host population might be realistic. Migration rates between compartments have been estimated to be small in HCMV samples from congenitally infected infants, Renzette et al., 2013. With our model we aim to represent the evolution of the persistent parasite population. This population is thought to be located primarily in CD34+ bone marrow cells (Dupont and Reeves, 2016). The bone marrow is distributed over several bones, this population structure could also be transferred to the persistent CMV population. Currently, no estimates of this population structure is available so that the simplest assumption of panmixia should be preferred.
- (viral reproduction) We model the evolution of a reproducing persistent viral population. On the real time scale the reproduction rate obviously is not constant. There are phases at which the immune system essentially silences viral reproduction and phases at which the virus reactivates. In phases of reactivation virus is produced and shed into bodily fluids to infect other hosts, but also cells are infected in which the virus eventually will persist. In our model tacitly the time for each host is rescaled accordingly to the speed of viral reproduction. There is increasing evidence, that HCMV reproduction is not restricted to early childhood, but continues at a lower frequency in (the immunocompetent) adulthood, especially in the eldery, see Boven et al., 2017; Huang et al., 2018. In particular, van Boven et al. hypothezise that “infectious reactivation in adults is an important driver of transmission of CMV”. For estimates of the viral reproduction rates see the separate point “parameter regimes”.
- (primary infection with a single type) On a long evolutionary timescale most hosts should be primary infected with HCMV via breast milk or in early childhood via saliva or urine, because in countries with a lower hygenic standard and/or high breast feeding rates the HCMV seroprevalence lies already at an age of xxx at xx%, see Adland et al., 2015 and references therein. The low virulence of HCMV (see also the point “relationship of the reinfection rate *r*_*N*_ and the parasite reproduction rate *g*_*N*_”) suggests that at primary infection most frequently only a single type can establish the infection in the new host. However, it has also been observed that during lactation several CMV genotypes have been transmitted to a preterm infant, see Hamprecht et al., 2003. The results about the maintenance of diversity in the setting with balancing selection also hold, if at a primary infection more than a single type is transmitted. Only the dynamical system driving the evolution of the host type frequencies might change in favour of a higher proportion of hosts being infected with both types. The case of neutral parasite reproduction we discuss in Remark 1 and Remark 2. We argue that the results of Section 3.1.1 and 3.1.2 also hold for a more general models of primary infection. The only exception is Case b) i) in Section 3.1.2 for primary infections at which many parasites found the persistent parasite population. However, in this case typically a host is hit by a host replacement before a reinfection event happens. For HCMV this scenario seems not to be relevant, since empirical studies indicate that diversity is accumulating over the life of a host, see also the point “reinfection”.
- (reinfection) It appears that HCMV can superinfect persistently infected hosts despite CMV-specific humoral and cellular immunity, see Hansen et al., 2010. Women who have been infected with CMV recently have been shown to be infected with a single CMV genotype only at the ORFs UL55, UL144, UL146 and UL09, see Murthy et al., 2011. On the other hand in the majority (15/16) of random samples from CMV-seropositive women (with the same demographic origin) with presumably past primary infections (as the women have been randomly sampled) several genotypes at the ORF UL55 and UL73 have been detected, see Novak et al., 2008. In Görzer et al., 2010 genotyping was performed for gB (UL55), gH (UL75) and UL10 for 36 immunocompetent patients during primary infection and in 14 patients experiencing HCMV reactivation/reinfection. In all cases only one gB-gH-UL10 genotype was detected in patients experiencing a primary infection, in contrast in 4 of 14 immunocompotent patients a mixed genotype infection was detected. The frequency of mixed infection thus differed significantly between primary and past infection (p-value =0.004, Fisher’s exact test). These findings indicate that also virions that are transmitted to a host at superinfection may persist in a host and therefore at least part of the diversity observed in hosts may have been introduced at (a/several) reinfection event(s).
- (balancing selection within hosts) In variant b) of the model we assume that both genotypes are actively maintained in host populations. Various mechanisms may lead to such an effect. For example HCMV exhibits a broad cell-tropism, HCMV particles can infect many different cell types of the human host, see Sinzger et al., 2008. The reproduction rate varies between different cell types and between HCMV samples. For example for virus entry intro fibroblasts only the complex gH/gL/gO is necessary, but for virus entry into epithelial cells both complexes gH/gL/UL128-UL131 and gH/gL/gO are necessary. In vivo reproduction in both cell types is important (Sinzger et al., 2008). Reproduction in epithelial cells is thought to be important for inter-host transmission, since these cells line the outer surfaces of organs and blood vessels and so virus can diffuse form these cells into bodily fluids. Reproduction in the ubiquitous fibroblasts provides the platform for efficient proliferation of the virus. Zhou et al. suggest that the different genotypes of gO might affect the ratio gH/gL/gO to gH/gL/UL128-131, see Zhou et al., 2013. In a host several gO genotypes might be maintained in the persistent viral population, so that both virus entry into epithelial as well as virus entry into fibroblasts may be tweaked. Another example are immune evasion strategies or mechanism for immune modulation. Different antibody responses exist against the antigens of genotypes gH1 and gH2 of the glycoprotein gH (UL75) and against two different antigenes of gB (UL55), see Urban et al., 1992; Meyer et al., 1992. Both genotypes may be controlled according to some predator-prey-dynamics with the immune system of the host on the one hand, on the other hand they may compete with each other due to a restricted amount of cells which can be latently/persistently infected.
- (parameter regimes) – (parasite population size *N* within hosts) Currently, there are no estimates of the effective size of the persistent viral population in the healthy host available. Only the population histories of congenitally infected infants, in which a latent phase has not been reached yet, have been inferred. The effective compartmental population sizes of the HCMV populations were estimated to be of order 1000, see Renzette et al., 2013. The actual number of virions infecting latently a host we estimate to be similarly high, to wit roughly between 2 · 10^3^ and 2 · 10^5^. We estimated the actual number of virions as follows: It is assumed that HCMV latently infects pluripotent CD34+ bone marrow progenitor cells (BMCs). In Slobedman and Mocarski, 1999 0.004-0.01 % of granulocyte colony-stimulating factor (GCS-F)-mobilized BMCs ^5^ have been estimated to be infected with HCMV and in this study HCMV-infected cells contained between 1 to 13 genomes. In Khaiboullina et al., 2004 0.1 ± 0.02% of CD 34+ BMCs have been estimated to be infected with HCMV in HCMV-seropositve healthy hosts. The number of HCMV genomes per infected cell have not been estimated in this study. The majority of CD 34+ cells are contained in the bone marrow. The estimated mean of the number of CD 34+ BMCs per kg body weight is 2.4 · 10^6^, see Körbling and Anderlini, 2001. Assuming a body weight of 70 kg for a standard individual, we estimate that the total count of HCMV virions contained in CD 34+ BCMs lies between 0.00004 · 70 · 2.4 · 10^6^ and 0.0001 · 13 · · 70 · 2.4 · 10^6^. – (host population size *M*_*N*_) The effective population size of the human population is estimated to be 10^4^, see Park, 2011. – (parasite reproduction rate *γ*_*N*_) The replication cycle of HCMV in HCMV-naive and HCMV-experienced hosts has been analyzed by Emery et al., 2002, see also the Review Griffiths et al., 2015. HCMV replication dynamics were investigated in a population of 30 liver transplant recipients with active HCMV replication. The mean growth rate of virus during infection of HCMV-naive individuals was 1.82 units/day (95% confidence interval (CI), 1.44– 2.56 units/day), corresponding to a viral doubling time of 0.38 units/day (9 h; 95% CI, 0.27– 0.48 units/day). Infection of patients with preexisting HCMV immunity was associated with a significantly slower viral growth rate (0.62 unit/day; 95% CI, 0.54– 0.71 unit/day) and a viral doubling time (1.12 days; 95% CI, 0.99– 1.25 days). Assuming that a typical host is infected for 50 years (by assuming that on an evolutionary time scale a typical host experiences an HCMV primary infection in early childhood) and the expected life time is about 50 years we arrive at an upper bound of (356/0.38) · 50 ≈ 48000 viral generations within a host. There is evidence, that in early childhood the virus replicates over a long period, Cannon et al., 2011. Assuming that in early childhood the virus replicates for two years (not necessarily at a stretch, within the first 5 years of infection), that is 365 · 2 days, we arrive at a lower bound of 365 · 2/1.12 ≈ 651 viral generations during this time period. In the eldery host the virus regularly spreads, see Stowe et al., 2007 in this study 57% of the investigated elderly hosts (which were older than 65 years old) were shedding CMV. On an evolutionary time scale these old hosts might not play a role. Therefore, for a lower bound on the number of viral generations we ignore this time period. Between early childhood and eldery the immune system is capable to control the virus in the healthy host pretty well. In healthy non-pregnant women the averaged shedding prevalence was estimated to be 4.7 %, see Huang et al., 2018. Hence, during this period we assume, that the virus is active each year at 0.047· 365≈ 18 days, which leads to roughly 9 viral generations per year. Hence, assuming again that a typical host is infected for 50 years, we arrive at a lower bound of 651 + 45· 18/1.12 = 1374 viral generations per host life. – (mutation rate *µ*_*N*_) The average mutation rate per site per replication cycle for HCMV has been estimated to be 2 · 10^−7^, see Renzette et al., 2011. In this study we are not interested in the mutation rate per site, but the mutation rate of entire reading frames, which genotype landscape clusters into a few types, see Puchhamer-Stöckl and Görzer, 2011 for an overview. As an example we consider the ORF UL75 encoding for the glycoprotein gH. This glycoprotein forms together with the glycoproteine gL (UL115) and either with the proteins encoded by UL128, UL130 and UL131 a pentameric complex or with the glycoprotein gO encoded by UL74 a trimeric complex, both of which are essential for the viral entry into epithelial/endothelial cells and fibroblasts, see Ryckman et al., 2008; Zhou et al., 2015; Ciferri et al., 2015. UL128, UL130 and UL131 are highly conserved regions, but the DNA sequences of UL74, UL75 and UL115 cluster into eight, two and four genotypes, respectively, see Puchhamer-Stöckl and Görzer, 2011 and the references therein. Two different antibody responses are triggered by the glycoprotein gH. The binding site of the neutralizing human antibodies react on two different variants of gH, which are distinguished by at least two amino-acid changes and an amino-acid deletion, see Urban et al., 1992. We separated the 132 consensus sequences ^6^ of low-passaged HCMV-samples provided by Sijmons et al., 2015 according to these amino-acid changes and the amino-acid deletion into two groups. according to this criterium in addition to the two amino-acid changes and the deletion further six amino-acid changes are separating the two genotypes. All eight amino-acid changes and the amino-acid deletion are located within a small N-terminal region of a length of 28 amino-acids. A high differentiation between the two groups of sequences is reflected in a high fixation index *F*_*ST*_ of 0.89, which has been calculated according to the formula by Hudson et al., 1992. Furthermore, the sequences of the two genotypes cluster phylogenetically into two groups, except for three sequences which coincide at the N-terminal region with the Merlin genotype, but carry further downstream patterns which are only present in the sequences of the AD169 genotype, see Supplementary Figure 1. In the literature sequences of the ORF UL75 are often separated into two groups, which are then called genotype gH1 and genotype gH2. However the criteria to be clustered into genotypes gH1 or genotype gH2 vary between publications. Here we use the 8 amino-acid changes and the amino-acid deletion. Epithelial/endothelial cells and fibroblasts are all cells types in which the virus can reproduce efficiently, see Sinzger et al., 2008 and from which virus is released into bodily fluids. As the ORF UL75 is strongly involved in virus entry into these cells, virus sampled from bodily fluids should carry an intact ORF UL75. In samples (from Sijmons et al., 2015) only the two genotypes gH1 and gH2 are detected (at least on the basis of a consensus sequence). Therefore we will assume in the subsequent analysis that only the two genotypes (which we distinguish by 8 amino-acid changes and one amino-acid deletion) allow a proper virus entry into these cells. As a consequence of our assumption all 8 amino-acid changes and the amino-acid deletion have to occur within a host life. If we assume in the following that the rate at which amino acids are deleted is the same as the mutation rate per site, the rate at which a change from genotype gH1 to genotype gH2 and vice-verse happens is (*g*_*N*_ · 2 · 10^−7^)^9^, because within a host life on average *g*_*N*_ viral reproduction events occur, the mutation rate per site was estimated to be 2 · 10^−7^ and all 8 amino-acid changes plus the amino-acid deletion have to occur. This leads to an upper estimate of (4.8 · 10^4^ · 2 · 10^−7^)^9^ ≈ 10^−18^ for the mutation rate *µ*_*N*_ per parasite, see the point “viral reproduction rate *g*_*N*_”. – (reinfection rates and the relationship of the reinfection rate *r*_*N*_ and the parasite reproduction rate *g*_*N*_) For simplicity, we do not assume, that the reinfection rate depends on the host state in our model. Currently, there is no data available, how the reinfection rate is influenced by diversity. We suggest that accumulating diversity (with respect to the whole genome) may antagonize the host’s immune system, and hence increases the reactivation rate and therefore increases also the reinfection rate. However, in this study we limit ourselves to a single locus and hence act on the assumption that the reinfection rate is homogeneous among all hosts. There have been used two different methods to assess the frequency of reinfection. In both cases longitudinal samples from patients were taken. In method a), see Bale et al., 1996, the samples (from the different time points) were sequenced at specific ORFs and the genotypes present in the different samples were compared. The appearance of a new genotype was considered as evidence for reinfection. In method b), see Ross et al., 2010, the reactivity to antibodies specific for certain genotypes of the ORFs UL55 and UL75 have been analyzed in time samples. The appearance of a new antibody specifity was considered as evidence for reinfection. Both methods have certain drawbacks. Using deep-sequencing methods it has been shown that genotypes are often present at very low frequencies only and that frequencies of genotypes vary strongly over time. At reactivation viral growth is presumably exponential. The virions which initiate the reactivation at the very beginning can grow most and the progeny of these few virions consitute the bulk of the viral population in bodily fluids from which virus is sampled. Therefore the frequencies of the genotypes measured in a single sample might not map well the frequencies of the genotypes in the persistent viral population and often may overrepresent a single genotype which is the type of the virion initiating the reactivation. Furthermore, in samples from different time points estimates of genotype frequencies may differ strongly, because the virions sampled may stem from different reactivations. Hence, with methods a) it well might be that the genotypes detected at the different time points were present in the host all the time. This would lead to an overestimation of the reinfection rate. A problem with using method b) is that a large proportion of hosts (roughly 30%) does not show any genotype-specific antibody reactivity. At least for ORF UL75 the reason cannot be that the hosts react to a different epitope, because in this region the population clusters only into two genotypes. In Ross et al., 2010 the authors claim that reactivity to strain specific epitopes was shown to persist on average for 21 months. Hence, it well might be that if a specific genotype is not reactivated the antibody response to that genotype decreases over time, although virions carrying this genotype still persist in the host. A lacking antibody response on the other hand may facilitate the reactivation of the virions with that genotype and renew the antibody response which then is detected in the samples. A reinfection event however must not have occurred to generate this response. This would lead to an overestimation of the reinfection rate with method b). In Ross et al., 2010 an annual reinfection rate has been estimated by first determining the proportions of hosts reinfected in a certain time interval and then dividing this number by the length of the time interval (in time units of years). Using this algorithm the reinfection rate may be underestimated, since hosts who are reinfected with genotypes they have already been infected with earlier may be not recognized as reinfected. This is especially likely if only a few genotypes are considered. If for example 20% of the CMV-infected persons are infected only with genotype gH1, 30% of the persons are infected with both genotypes gH1 and gH2 and 50% of the persons are infected only with genotype gH2, a person (so far only infected with genotype gH1) is only in roughly 30·0.5 % + 50% of all reinfections infected with genotype gH2 (if one assumes that a host infected with both genotypes transmits in 50% of the cases genotype gH1 and in 50% of the cases genotype gH2). By determining the proportion of the different genotypes in the population and transmission probabilities of the different genotypes from host infected with several genotypes, one should be able to estimate the reinfection rate quite well also when considering only a few (or a single) ORF (of course provided that one is able to distinguish the occurrence of a new genotype from the reactivation of an already present genotype). With method a) in 7 (19%) of 37 young children attending day-cares evidence for reinfection has been shown within a time frame of sampling between 11 weeks to 38 months with a median of 11 months, see Bale et al., 1996. In Ross et al., 2010 205 CMV seropositve women and in Yamamoto et al., 2010 149 CMV pregnant women (40 of these with congenitally infected infants and 109 with uninfected newborns) were followed prospectively with method b). In the first study an annual reinfection rate of 10% has been estimated and in the second study an annual reinfection rate of 34% for mothers with congenitally infected infants and an annual rate of 9% has been estimated for mothers with uninfected newborns. A similar primary infection rate has been estimated in a study population with the same geographical origin, see Colugnati et al., 2007. It is not clear, why the primary infection rate is not higher than the reinfection rate, because preexisting immune response should prevent at least partially recurrent infections. We assume that the reinfection rate is smaller than the viral reproduction rate. The reasoning is that before a host can reinfect another host the virus has to replicate first within this host. Furthermore the virulence of HCMV is pretty low. Transmission over breast milk is relatively ineffective. Nearly every (96 %) HCMV-positive mother reactivates HCMV when nursing, see Hamprecht et al., 2001. However, transmission often (in approximately 80% of the cases) results only in local oral infections not leading to a persistent infection with HCMV, see Mayer et al., 2017. In the study of Hamprecht et al., 2001 a mother-to-infant transmission rate of 37% was estimated and viral DNA or infectious virus was detectable in the infant with a mean of 42 days (95% CI [28,69]) after the estimated onset of transmission risk in the mother. Similarly, the transmission over saliva requires time to be effective. The annual seroconversion rate of women with infected children was estimated to be 30% and for mothers with infected children younger than 20 months of age the interval between identification of her child’s infection and maternal infection ranged from 1 to 26 months (8 +/-6 (SD) months), see Adler, 1991b.

#### 3.2.2 HCMV data

##### Materials

Apart from data from the literature we base our study on genotype data for the open reading frame UL75 from 17 immunocompetent patients which were collected retrospectively for a previous study (Görzer et al., 2010) with ethical approval of the review board of the Medical University of Vienna (EK Nr: 202/2009).

From all patients serum, saliva, and urine samples were sent to the Center for Virology, Medical University of Vienna between 2001 and 2008 and tested for HCMV-specific IgG antibody avidity by a previously described avidity assay (Steininger et al., 2004) and for HCMV-DNA since they had clinical symptoms as fever, hepatosplenomegaly, elevated transaminases, lymphadenopathy, and/or suspected HCMV infection. Twenty-one out of 84 patients had a high IgG antibody avidity indicative for a past HCMV infection, see Prince and Lapé-Nixon, 2014. In each sample the genotypes gH1 and gH2 of the region UL75 were detected using genotype-specic qPCR. The detection threshold for each of the two genotype gH1 and gH2 was 100 copies/ml, see Görzer et al., 2008. We excluded 4 from the 21 patients for which the number of viral particles was below 1000 copies/ml, because in this case a type present at frequency 0.1 is likely not to be detected in the sample. Patients for which the frequency of one of the two types was below 0.1 is treated as purely infected The youngest patient was 2 years old, the oldest 84.1 years, the mean age was 55.15 years with a standard deviation of 25.35 years. For two patients saliva samples were taken, for 15 patients serum samples (see Supplementary Table 1).

We excluded the 63 hosts with a low HCMV IgG antibody avidity, indicating an ongoing primary infection with HCMV. These hosts were signicantly more frequently infected only with a single genotype. In 35 of these 63 immunocompetent patients (from the same hospital, the same time interval for sample collection and with the same symptoms as for the 17 patients) an HCMV-DNA load of at least 1000 copies/ml was detected. In 34 patients a single gH genotype was detected and both genotypes were found in a single host with the minor type at frequency 0.013 (see Supplementary Table 2). The number of hosts infected with both types is in the hosts with low IgG avidity signicantly lower than in the other 17 hosts with high IgG avidity (p-value = 0.0036 using Mann-Whitney’s U-test)). Since also in random samples from immunocompetent children a higher proportion of children was infected with both genotypes, see Paradowska et al., 2014, we excluded the samples with low IgG avidity from the study.

#### 3.2.3 Coexistence of both parasite types due to mutations at the example of HCMV

We are interested in the probability to detect in a sample of parasites two different parasite types. We base again our analysis on the data available for the ORF UL75. We already discussed in the point “mutation rate *µ*_*N*_” that on the host time scale we estimate the rate at which a virion changes its type from genotype gH1 to genotype gH2 by 10^−18^, if we assume that only the genotypes gH1 and gH2 (and not intermediate variants of them) can be propagated further.

According to the results from Section 3.1.1 the total tree length of a sample of size *n* taken from the entire parasite population is at most of order log(*n*) min {*M, MN/r*_*N*_} on the host time scale.

Furthermore, according to our assumptions the most recent common ancestor of the sample is either of genotype gH1 or gH2. So both genotypes are only detected in the sample, if on the genealogy of the sample falls at least one change from genotype gH1 to gH2 or vice-versa. The probability that this happens can be estimated from above by log(*n*) · *M* · 10^−18^. With *n* = 132 and an estimate of the effective host population size of 10^4^, we arrive at an upper estimate of 5 · 10^−14^ to observe both genotypes simultaneously in the population. To observe even more genotypes is even less likely.

If intermediate types can be propagated as well the detection of both genotypes in a sample is more likely. However, in this case also intermediate variants should be observed with high probability, which is not the case.

One might argue that just by chance the genotypes gH1 and gH2 are present at the observed frequencies in the population. This scenario we discuss in the next section.

#### 3.2.4 Diversity due to standing genetic variation at the example of HCMV

The proportion of hosts infected with both genotypes is in our sample 3/17= 0.176 (with a 95%-confidence interval of (1/17,5/17)). These estimates accord with the estimates from Paradowska et al., 2014, were the frequency of neonates infected with both types were 14.3%, of infants was 23.7% and of adults was 36.4%. In this paper neonates were congenitally infected, infants were infected postnatally (or by an unproven congenitally CMV infection) and adults were HIV patients or kidney recipients. Since these adults are immunocompromised it is likely that they are more susceptible for reinfections, which may explain the higher rate (in comparison to our estimate) of mixed infections in adults.

Furthermore, three patients (17.6%) are purely infected with type gH1 and 11 (64.7%) patients only with type gH2. In Paradowska et al., 2014 the proportion of hosts only infected with type gH2 was also in all patients groups larger than the proportion of hosts only infected with type gH1. The discrepancy was largest in adults (40.5% to 45.2% in newborns, 33.3% to 43% in infants and 25.4% to 38.2% in adults).

The average frequency of type gH1 is 0.251. Hence we estimate 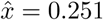.

In the neutral case the highest proportion of hosts infected with both types is expected, if *r*_*N*_ *≫ N* and *r*_*N*_ *≈ cg*_*N*_ for some *c >* 0. According to the results from Section 3.1.2 in this case the proportion of hosts infected with both types is Beta distributed with parameters 0.251*c* and 0.251(1 *- c*). As pointed out already only hosts with a frequency of type gH1 between 0.1 and 0.9 are treated as mixed (i.e. infected with both types). Hence, to estimate *c* we find the maximal *c* such that 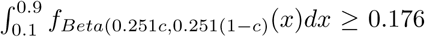 (and to determine the confidence interval we find the maximal *c* such that the integral *≥*1/17 and *≥*5/17, respectively). This leads to an estimate of *c* = 0.26 (with a 95% confidence interval of (1/15, 0.49)).

That means that *r*_*N*_ ≈ 0.26*g*_*N*_. So, in the neutral case roughly every fourth viral generation a reinfection event should happen.

Assuming that within a host life on average 1347 viral generations occur, we arrive at an average number of 350 reinfections per host life, that is 7 reinfection per year, which is 70 times higher than the estimated reinfection rates of Ross et al., 2010.

Note that these estimates also hold true in more general models for primary infection as we discussed under the point “primary infection with a single type”.

## 4 Discussion

In this study we examined the diversity patterns in parasite populations which are capable for persistence and reinfection. In Pokalyuk and Wakolbinger, 2018 a model for the evolution of a population of parasites has been introduced. In this model parasites are distributed over a large number *M* of (infected) hosts. Each host is infected with a large number *N* of parasites and there exist two parasite types. Evolution is driven by host replacement, reinfection and parasite reproduction. In Pokalyuk and Wakolbinger, 2018 it is assumed that parasite reproduction is driven by balancing selection, that is there is a tendency that both parasite types are maintained in a host (until a host dies). In this case it has been shown in Pokalyuk and Wakolbinger, 2018 that persistence and reinfection are (in certain parameter regimes) effective mechanisms two maintain diversity in a parasite population.

Mechanisms allowing the maintenance of diversity (especially in coding regions) in a parasite population might be essential for its survival. Only if the parasite population is sufficiently diverse it can manage to survive in its host, otherwise the host can adapt its host defense to easily against the parasite. We propose that persistence and reinfection might be essential mechanisms enabling a parasite to accumulate sufficient diversity within hosts without generating diversity constantly de novo by mutation.

Certain diversity patterns observed in HCMV samples motivated the model of Pokalyuk and Wakolbinger, 2018. In coding regions on the HCMV genome the observed diversity is rather high. The HCMV genome possess roughly 171 canonical, protein coding ORF, see Dolan et al., 2004; Gatherer et al., 2011. At least 23 of these cluster into more than one genotype, see Puchhamer-Stöckl and Görzer, 2011, that is more than 13 %^7^. Furthermore, when these ORFs have been analyzed further, a large proportion of hosts was infected with several genotypes. These observations seem to contradict neutral evolution.

In this study we analysed the neutral counterpart of the model, that is with neutral parasite reproduction within hosts, to lay the basis for a fit of the neutral model to data. In the neutral case reinfection may allow the spread of parasite types in a population, however reinfection and persistence do not enhance the long-term coexistence of the two types in the parasite population - fundamentally different from the model with balancing selection. We give asymptotic upper bounds for the length of the total genealogy of a random sample taken from the parasite population which allows to estimate the probability of coexistence of two types. Furthermore, we analyzed the genealogy of a sample taken in a single host, which enables us to determine the asymptotic distribution of the proportions of two parasite types in single hosts (given both types coexist in the population). It turns out that these asymptotic distributions are parametrized by the relationship of the parasite reproduction rate *g*_*N*_ and reinfection rate *r*_*N*_.

We used these results to evaluate the relevance of the neutral scenario for HCMV. On the basis of the literature and genotyping data at the ORF UL75 (clustering into two genotypes) from 17 immunocompetent patients we infer that both genotypes should coexist in the HCMV population at most with probability 5 · 10^−14^. This estimate is based on the number of amino-acid changes and deletions which distinguish the two genotypes (for at least 132 consensus sequences) and the assumption that intermediate variants are not propagated between hosts. Further empirical studies are necessary to verify this separation into genotypes and the hypothesis of transmissability. Genotypes of other ORFs might be distinguished by less amino-acids and therefore coexistence might be more likely just due to mutation. Nevertheless this estimate conflicts with observation that at least 13% of all HCMV ORFs cluster into several genotypes.

Given that two genotypes coexist in the parasite population at the ORF UL75 we inferred (from the 17 patient samples) that the viral reproduction rate is (roughly) only 4 times larger than the reinfection rate. Based on estimates of the viral reproduction rate inferred from the literature this would mean that every year a host is reinfected at least 7 times. This estimate is 70 times larger than current estimates of the reinfection rate.

A larger set of ORFs should be analyzed to reveal a more precise picture. However, our estimates provide a first indication that other (non-neutral) scenarios might mesh better with the observed diversity patterns. Our current data set is too small to fit the data to the dynamical system driving the evolution of the parasite model with balancing selection. However, the parameter regime analyzed in Pokalyuk and Wakolbinger, 2018 seems to be applicable for HCMV. In particular, in the model with balancing selection two parasite types are likely to be observed also for small(er) reinfection rates. The major constraints on the model are that the selection strength *s*_*N*_ = *N* ^*−b*^ for some 0 *< b <* 1 and scales proportionally inverse to *r*_*N*_. In addition it is required that *g*_*N*_ ≪ *N*^3*b*+*E*^. ^8^ We estimated *g*_*N*_ to be bounded from below by roughly 1000. The size of the persistent parasite populations within hosts has not been quantified so far, we argued that it might be of the order of 1000. As an example consider *b* = 0.2 and *N* = 1000, then *N* ^3*b*^ ≈ 63 and for *N* = 10000 we have *N* ^3*b*^ ≈ 250. Both values are much smaller than *g*_*N*_. Furthermore 1*/s*_*N*_ ≈ 4 for *N* = 1000 and ≈ 6 for *N* = 10000, which is roughly on the scale of the reinfection rates which have been estimated in the literature.

Several extensions of the model considered here are relevant. In the parameter regimes suitable for HCMV we discussed already here that our estimates concerning HCMV hold also for more general models of host replacement, to wit that at primary infection several types can be transmitted. In the case with balancing selection a more general model of primary infection may just lead to a larger proportion of hosts infected with both parasite types. Regarding persistence and reinfection as mechanisms which allow in combination with balancing selection the accumulation of diversity within hosts (and in this manner contributing to the long-term coexistence of HCMV with its host), models treating several loci (all allowing for several genotypes) simultaneously are particularly relevant extensions.

## Acknowledgements

CP thanks Anton Wakolbinger for fruitful discussions. CP was supported by the DFG priority programme 1590.

## Supplementary Material

**Supplementary Figure 1:**
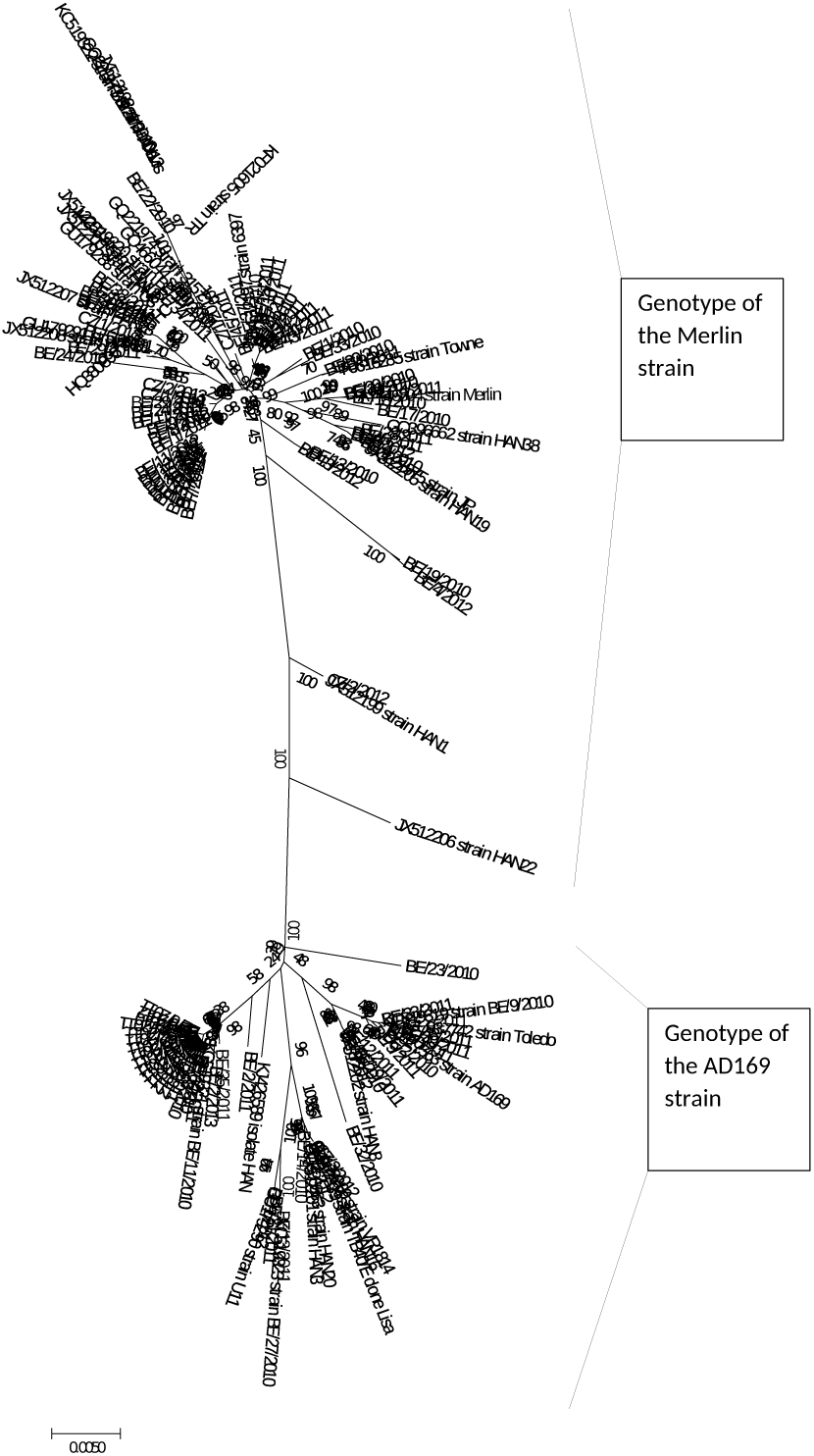
The evolutionary history was inferred using the Neighbor-Joining method (Saitou and Nei, 1987). The optimal tree with the sum of branch length = 0.24032069 is shown. The percentage of replicate trees in which the associated taxa clustered together in the bootstrap test (500 replicates) are shown next to the branches (Felsenstein, 1985). The tree is drawn to scale, with branch lengths in the same units as those of the evolutionary distances used to infer the phylogenetic tree. The evolutionary distances were computed using the p-distance method (Nei and Kumar, 2000) and are in the units of the number of base differences per site. The analysis involved the 132 nucleotide sequences from Sijmons et al., 2015. Codon positions included were 1st+2nd+3rd+Noncoding. All positions containing gaps and missing data were eliminated. There were a total of 2226 positions in the final dataset. Evolutionary analyses were conducted in MEGA7 (Kumar et al., 2016).

**Supplementary Table 1.**
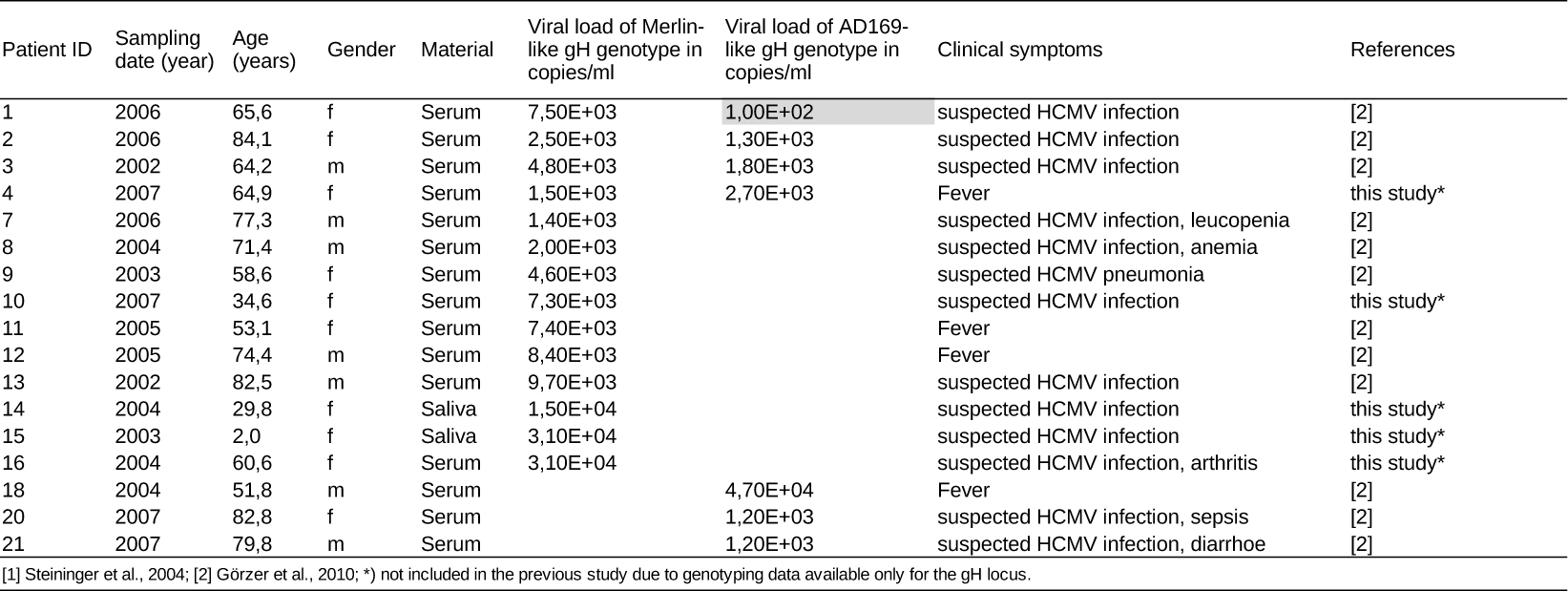
Patients with past HCMV infection as proven by high HCMV-specific IgG antibodies

**Supplementary Table 2.**
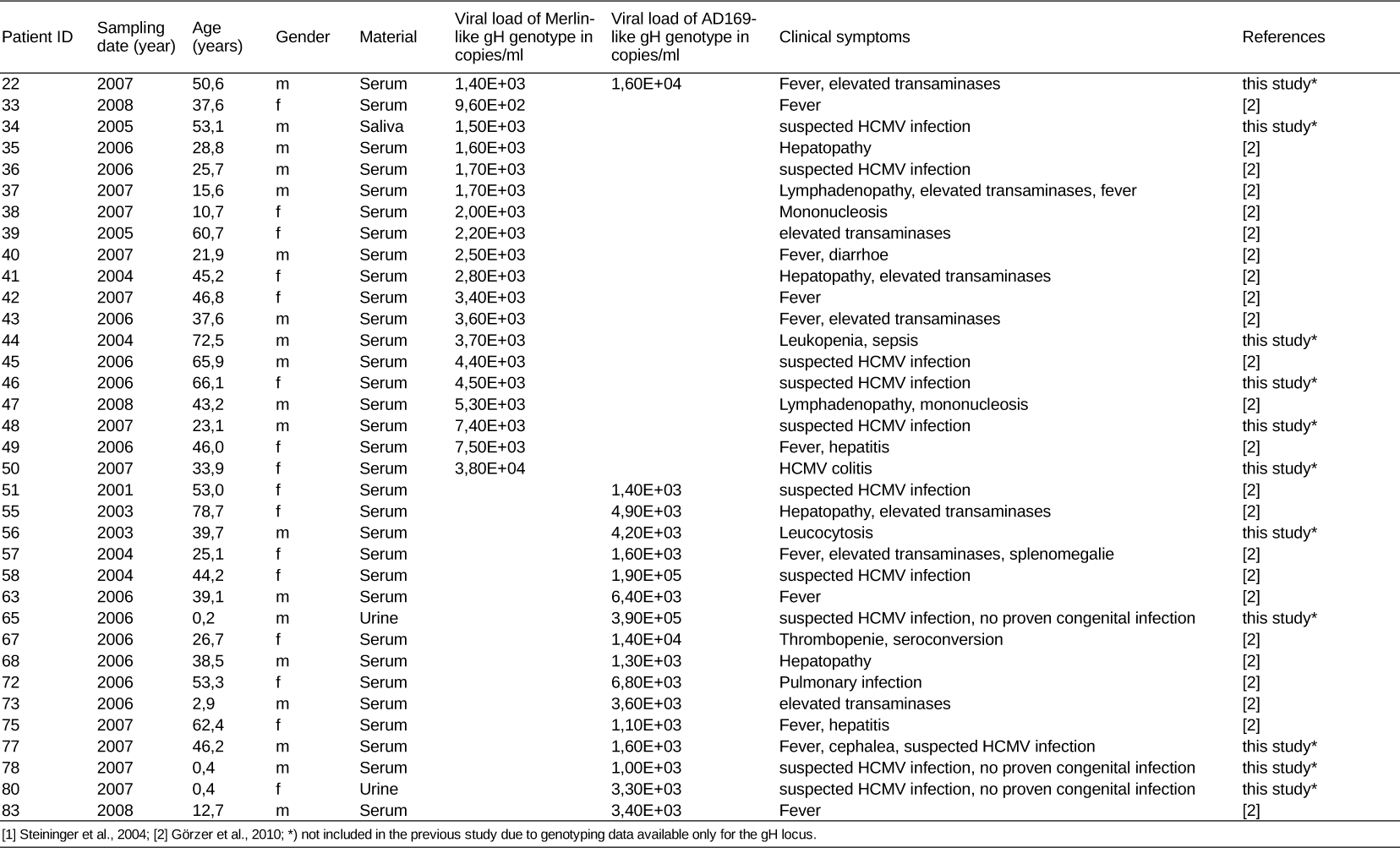
Patients with primary HCMV infection as proven by low HCMV-specific IgG antibodies and/or seroconversion.

## Footnotes

1 as the resulting distribution is again continuous and memoryless

2 that is viral mechanisms to infer with the immune response of the host

3 that is allowing HCMV to replicate in different cell types of the host

4 these cells are thought to the main site of the persistent HCMV population

5 CD34+ BMCs are mobilized with GCS-F to migrate to the peripheral blood where they can be sampled more easily. There is no indication that the proportion of CD34+ BMCs infected with HCMV is altered by this treatment

6 A consensus sequence is determined as follows: The HCMV DNA from e.g. a urine sample of a HCMV-positive patient is sequenced and the reads are aligned to a reference sequence. Then the consensus sequence is built from this alignment by choosing the most frequent variant at each site.

7 In Stern-Ginossar et al., 2012 have been annotated in total 751 protein coding ORFs. However most of the novel identified ORFs are very short (245 are less than 20 codons long and 239 ORFs are between 21-80 codons long), so that most of the underlying fitness landscape presumably do not allow for several genotypes. Furthermore these codons are found upstream of longer ORFs. 120 novel identified ORF were longer than 120 codons long. These are primarily ORFs contain slice junctions or alternative 5’ ends of previous annotations. Hence the observed diversity might coincide for these ORFs with that of the previously annotated ORFs.

8 The upper bounds on the mutation rate are not essential here, because in this case only a larger proportion of hosts infected with both genotypes is expected.

